# Voluntary Dissociation of Motor Unit Activity in the Vastii Muscles

**DOI:** 10.1101/2025.10.20.683602

**Authors:** Daniel Haller, Finja Beerman, Raul C. Sîmpetru, Luca Hofbeck, Roger M. Enoka, Alessandro Del Vecchio

**Affiliations:** Neuromuscular Physiology and Neural Interfacing Laboratory, Friedrich-Alexander-Universität Erlangen-Nürnberg, Erlangen, Germany; Traumatology and Orthopaedics, Universitätsklinikum Erlangen, Germany; University of Colorado Boulder, USA

## Abstract

The CNS coordinates movement through consistent activation patterns across muscles and motor units, suggesting the presence of a relatively fixed and high-dimensional number of neural constraints on voluntary actions. In the human quadriceps, the vastus medialis (VM) and vastus lateralis (VL) control the knee extensor torque and are considered a synergistic pair largely activated by shared neural inputs. However, some evidence suggests that these muscles, or even subregions within them, can be controlled independently. In this study, we investigated whether humans can dissociate neural input to VM and VL during isometric contractions. Ten participants received real-time feedback from multiple intramuscular EMG electrodes that targeted different regions of the VM and VL while attempting to activate each muscle or sub-regions selectively. We found that nine out of ten subjects were able to clearly separate VM and VL activity based on the intramuscular EMG feedback. However, motor unit decomposition from the intramuscular EMGs revealed that selective recruitment of a unique set of motor units was possible only within the proximal region of VM. In contrast, VL and distal VM showed highly correlated activation, indicating tight functional coupling. Correlation analyses confirmed that the proximal VM exhibited distinct activation profiles compared with both distal VM and VL, supporting the existence of compartmentalized control within VM. These findings demonstrate that it is possible to dissociate the activation of motor units within this synergistic muscle group during low-force isometric contractions.

## INTRODUCTION

The central nervous system (CNS) simplifies the control of movement by distributing common synaptic inputs to groups of motor neurons, organizing them into functional units often referred to as muscle synergies. This concept, rooted in the principle of dimensionality reduction, proposes that the CNS manages complex musculoskeletal systems by projecting a limited number of neural commands onto multiple muscles or motor units, enabling economical and coordinated motor behavior (d’Avella et al., 2006; Dernoncourt et al., 2024; Ivanenko et al., 2004; Laine et al., 2015; Luca and Erim, 1994; Madarshahian et al., 2021; Negro, Yavuz, Farina, 2016).

The prevailing view within the quadriceps muscle group has long been that the vastus medialis (VM) and vastus lateralis (VL) predominantly share neural input (Laine et al., 2015; Mellor and Hodges, 2005; Rossato et al., 2024). Consequently, VM and VL are typically considered a synergistic pair that acts collectively to produce a knee extensor torque and to maintain patellar stability (Laine et al., 2015; Powers, 2000). Such shared input limits the degrees of freedom and available motor-control strategies (Desmedt and Godaux, 1978; Desmedt JE, 1977; Marshall et al., 2022; Milner-Brown et al., 1973; Rossato et al., 2024).

However, studies that have leveraged high-density surface electromyography (HDsEMG) and motor unit decomposition during isometric tasks have revealed that there is some flexibility in the modulation of discharge rates of individual motor units in VM and VL (Del Vecchio et al., 2023; Levine et al., 2022; Rossato et al., 2024; Sandercock et al., 2018). By applying analytical techniques such as factor analysis (FA) and principal component analysis (PCA) to extract synergies (d’Avella et al., 2003; Tresch et al., 2006), distinct neural inputs to subsets of motor units within VM and VL have been identified. Despite these advances, it remains largely unknown whether humans can volitionally dissociate neural inputs to synergistic muscles that are functionally unified by the same tendon, such as the vastii muscles.

A key theoretical framework that has emerged in recent years is the concept of motor unit modes (Del Vecchio et al., 2023), which comprise groups of motor units that tend to be coactivated by common synaptic input across multiple tasks (Weinman et al., 2024). Although analyses at the population level have shown evidence of multiple neural modes, it remains unknown if human individuals can intentionally control these separate inputs in real-time tasks. To address this gap in knowledge, we combined intramuscular EMG recordings with real-time feedback of root-mean-square (RMS) amplitude and motor unit activity while participants explored voluntary strategies to activate different regions of VL and VM. We hypothesized that it would be possible to modulate neural inputs across tasks by recruiting distinct sets of motor units.

Our results demonstrate that most participants were able to dissociate activation of the proximal portion of the VM from the remaining synergistic muscle regions, including the distal VM and the VL. In a subset of participants, full dissociation between VM and VL was also achieved. These findings suggest that region-specific modulation of neural input is possible even within synergistic muscles, but this capability appears constrained by neuromechanical factors that are likely shaped by the coupling between muscle compartments. This appears to be the first study to combine intramuscular EMG with real-time feedback to show that humans can selectively recruit distinct sets of motor units within the quadriceps. These results reveal a previously underappreciated capacity for selective neuromuscular control, especially within proximal regions of muscles traditionally considered functionally unified.

## Methods

### Study Design

This study evaluated the ability of participants to modulate the neural input to the VM and VL, building on previous findings from HDsEMG and factor analysis of motor unit discharge rates (Del Vecchio et al., 2023; Levine et al., 2022). Each participant completed a single 30-min experimental session. Rather than following a rigid protocol, participants were encouraged to explore different voluntary strategies to achieve selective activation of either the VM or VL while maintaining low activation levels to facilitate accurate motor unit decomposition. All tasks were performed in a seated position on an isokinetic dynamometer, with the instrumented leg attached to the lever arm (Figure 1D).

**Figure 1.**
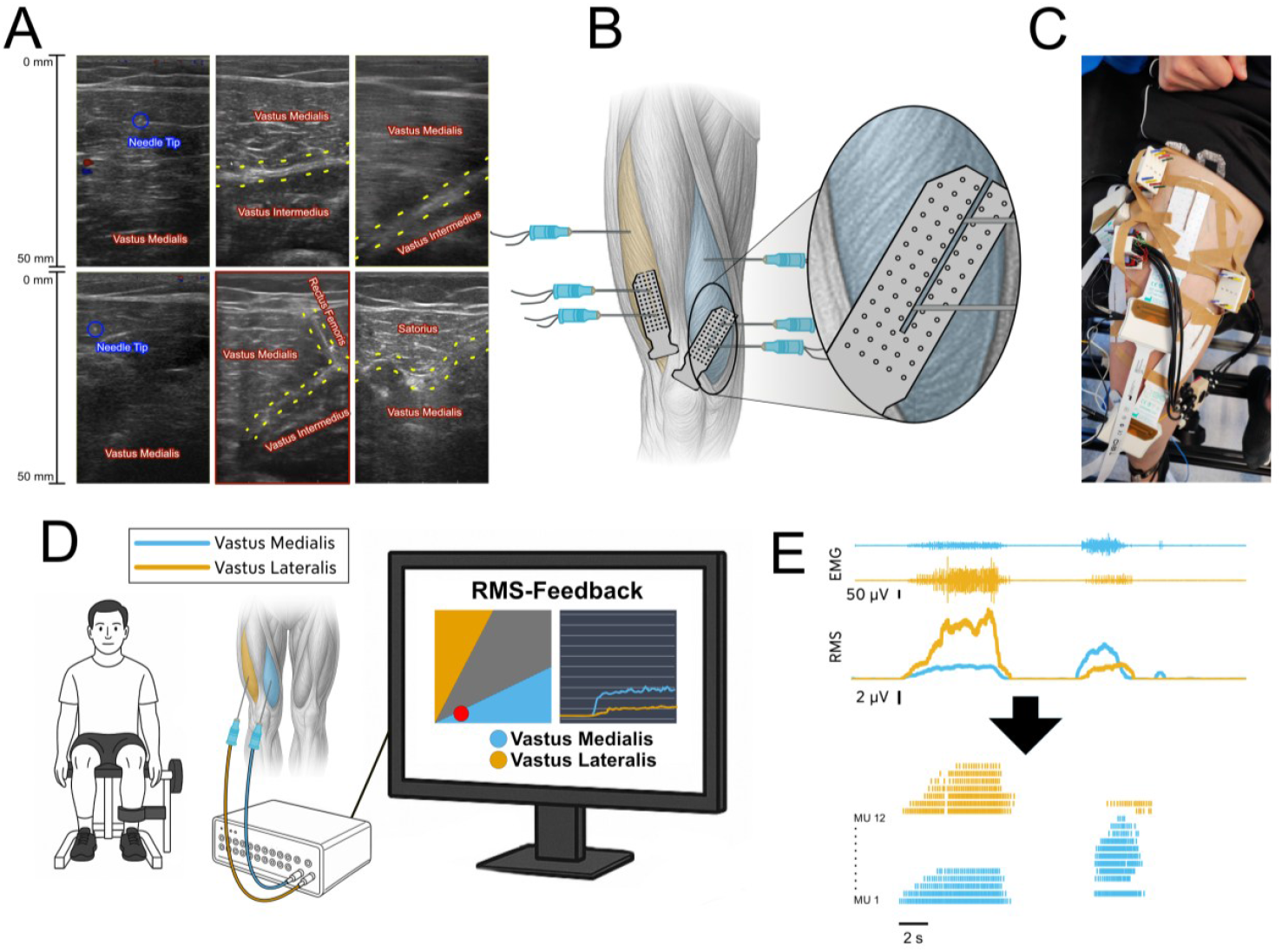
**(A) Anatomical visualization using ultrasound imaging** Six ultrasound images illustrate the anatomical identification and electrode placement within the quadriceps. The two images on the left display the needle tip within the vastus medialis (VM) during insertion, confirming intramuscular electrode position. The four images on the right depict transverse views of the quadriceps musculature at different depths and locations, highlighting clear separation between the VM, VL, and rectus femoris muscles. Muscle borders are manually outlined, and muscle labels are overlaid to validate targeted electrode placement. **(B) Electrode placement for intramuscular and surface EMG recordings** The schematic illustrates the placement of recording electrodes in the right quadriceps. The VL region is highlighted in yellow, and the VM in blue. Three intramuscular electrodes were inserted into each muscle (shown as needle symbols), with two electrodes positioned distally and one proximally in VM. HDsEMG grids were placed over the distal VM and VL regions. In VM, a small gap was cut into the grid to allow placement of the underlying intramuscular electrodes. This configuration ensured simultaneous recording of spatial surface signals and deeper intramuscular activity. The proximal intramuscular electrode in VM was always positioned outside the grid area to capture regional differences in muscle activation. **(C) Exemplary Final Electrode Setup on a leg** This photograph shows the complete electrode configuration on a participant’s leg. All intramuscular and surface EMG electrodes are visible, including grids over the VM and VL, as well as one placed on the rectus femoris. While the rectus femoris grid was applied during the experiment, data from this muscle were not analyzed in the present study. The image provides a practical overview of the real-world implementation of the setup and demonstrates the location of the electrodes across quadriceps muscles. **(D) Schematic representation of the experimental setup** This schematic illustrates the experimental setup during data collection. The participant was seated on an isokinetic dynamometer with the instrumented leg attached to the lever arm to ensure isometric conditions and prevent unintended movement. The screen in front of the participant displays the real-time EMG feedback used during the task. Two visualizations are shown: a 2D scatter plot of VM versus VL RMS amplitudes, and a time-series plot of RMS activity. This setup enabled participants to explore selective activation of VM and VL under controlled biomechanical conditions. **(E) Example of resulting processed signal** This panel illustrates EMG recordings and their associated processing steps for one channel each from VM (blue) and VL (yellow). The top trace shows the raw intramuscular EMG signals during alternating VM- and VL-focused tasks. Below, the RMS amplitude envelopes for each muscle are shown. The bottom trace displays the spike trains of identified motor units obtained through offline decomposition. Each row represents a single motor unit, with vertical ticks marking discharge times.

In this study, we defined a “VM-focused task” as any effort in which participants actively attempted to selectively increase activation in the VM while minimizing activation in the VL. Conversely, a “VL-focused task” refers to attempts to selectively activate the VL while minimizing activation in the VM. To help participants understand the task, we explained that they should attempt to contract one muscle more than the other by using internal cues such as focused attention or intentional muscle tension, visual cues by observing changes in the shape or movement of the muscle belly, and tactile cues by lightly touching the muscle to feel its contraction.

Throughout the session, participants received real-time feedback based on the RMS amplitude of intramuscular EMG signals, displayed either as time-series plots or as two-dimensional plots of VM and VL activity. Participants were told that these signals corresponded to the activity level of each individual muscle and were instructed to maintain low activation levels. Our primary outcomes of the study were 1) selective activation of VM or VL during real-time tasks, and 2) the identification of motor unit activity during VM- and VL-focused tasks, assessed via offline decomposition of intramuscular and surface EMG signals.

### Participants

Ten healthy individuals (mean age 29 ± 5.4 years; 6 males, 4 females) took part in the study. All were free from neurological or musculoskeletal disorders affecting the lower limbs. Written informed consent was obtained from all subjects prior to testing. The study protocol was approved by the ethics committee of the Friedrich-Alexander University Erlangen-Nürnberg (ethics vote 25-37_2-S) and was conducted in accordance with the Declaration of Helsinki.

### Experimental Protocol

Fine-wire intramuscular electrodes were inserted into both VM and VL under sterile conditions by trained medical personnel using standard anatomical landmarks and guided ultrasound imaging. The VM insertion was guided by identifying the medial border of the patella and tracing the muscle belly proximally along the medial aspect of the thigh. The VL landmarks included the lateral border of the patella and the midpoint between the greater trochanter and the lateral femoral condyle, following the lateral contour of the thigh. Electrode placement was verified using ultrasound imaging (Figure 1A). Although the fine-wire needle was often difficult to visualize, the tip could typically be identified, particularly when the needle was gently manipulated during insertion. Muscle borders were clearly visible. As an example, Figure 1A shows the borders between VM, vastus intermedius, and rectus femoris. This approach ensured that the electrode remained within the intended muscle belly and did not penetrate adjacent muscles. Visualization ensured that cross-talk from neighboring muscles was minimized and that recordings reflected true intramuscular activity from the targeted region. In each muscle, three electrodes were inserted to capture regional differences in activation (Figure 1B): two were positioned distally and one proximally.

To complement intramuscular recordings, high-density surface EMG (HDsEMG) grids were positioned over the skin above the distal electrode sites in both VM and VL. In VM, a small cutout in the grid allowed insertion of the underlying wires (Figure 1B, C). This configuration enabled simultaneous spatial mapping of surface and intramuscular motor unit action potentials.

Participants remained seated throughout the experiment in an isometric dynamometer setup. The instrumented leg was attached to the lever arm to prevent unintended movement. Participants were free to adjust the knee flexion angle and the rotation of the leg within the seated position to find a posture that best enabled selective activation of VM or VL. These adjustments were permitted during the exploration phase and between trials. The exploration phase lasted approximately five minutes at the beginning of the session. During this time, participants were encouraged to freely test different contraction strategies and postural adjustments while observing the EMG feedback. This allowed them to develop an intuitive understanding of how changes in effort, posture, or muscle engagement translated into visual feedback, helping them gain a sense of control over the system before beginning formal trials.

It was crucial that posture was held constant between VM- and VL-focused tasks within the same trial to ensure that motor unit shapes and decomposition accuracy were not affected by changes in joint angle and electrode geometry. While minor involuntary adjustments were not eliminated, all data used for analysis were visually monitored during acquisition to ensure a consistent posture across conditions.

Participants were instructed to focus on contracting either VM or VL individually (VM focused vs. VL focused task) using internal cues or subtle postural adjustments (e.g., slight changes in leg rotation or muscle pre-tension) to identify a strategy that allowed for selective activation of the target muscle. This exploratory design coupled with a real-time, zero-lag, highly intuitive interface of intramuscular EMG amplitude from VL and VM activity ensured that each participant adopted personalized and intuitive strategies for achieving selective activation, with the goal of recruiting unique sets of motor unit in either VM or VL. No external force targets were imposed, and participants were not required to produce measurable force output, as the primary goal was selective muscle activation rather than torque production. They were also instructed to include a brief relaxation period between tasks to ensure full derecruitment of motor units, thereby minimizing a potential influence of persistent inward currents (Beauchamp et al., 2025).

Participants were given approximately 30 minutes to explore voluntary control strategies and achieve the goal of selectively activating VM and VL. Rest periods were provided as needed to minimize reductions in force capacity. The task was considered successfully completed when participants demonstrated the ability to repeatedly and intentionally alternate between VM- and VL-focused activation, as reflected in the real-time EMG feedback. Trials were stopped when participants showed consistent and voluntary modulation of muscle activity, enabling clear shifts in feedback signals between the two tasks. This repeatable control pattern confirmed that participants had found a strategy to recruit each targeted muscle with minimal coactivation.

### Force-Direction Recording and Analysis

To complement the EMG recordings, we also measured three-dimensional force vectors during the selective activation tasks. The instrumented leg was fixed to the lever arm of the isokinetic dynamometer, constraining motion in the lateral (x-axis), upward (y-axis) and forward (z-axis) directions. To stabilize the foot, a small support platform was added. Because this platform was not mechanically connected to the dynamometer’s transducer, the absolute vertical force component was underestimated. However, the upward force was still partially captured by the sensor, making the vertical contribution to the task clearly visible in the recordings.

The force vectors were extracted under VM-focused (V_VM) and VL–focused (V_VL) conditions. To examine systematic differences between these conditions, we computed difference vectors ΔV = V_VL − V_VM for each trial. Principal component analysis (PCA) was then applied to ΔV across participants. The first principal component (PC1) explained the majority of the variance and its loadings on the three axes (x, y, z) identified the dominant directional component contributing to between-condition differences.

### Control Ramp Contractions

To assess whether the compartment-specific activation observed during EMG feedback tasks also emerged under conventional loading conditions, participants performed additional ramp contractions without EMG feedback. These control tasks were completed in the same isokinetic dynamometer, with all EMG electrodes and a 45-degree knee angle. Participants were instructed to perform two linear ramp contractions: one reaching 15% and one reaching 30% of maximal voluntary contraction (MVC) force, with a plateau phase of 30 seconds. Each task was repeated three times.

Force trajectories were displayed on a monitor, and participants were asked to follow them as smoothly as possible. No instructions regarding selective muscle activation were given. EMG signals were recorded from all intramuscular and surface channels as in the main experiment. For each ramp trial, root-mean-square (RMS) signals were computed for proximal and distal VM and VL channels. Pairwise Pearson correlation coefficients were calculated across these regions using the ramp-up phase to quantify the degree of co-modulation under default contraction strategies.

### EMG feedback

The feedback interface was implemented in a custom-written Python script and displayed either a two-dimensional scatter plot with VM-RMS amplitude plotted on one axis and VL-RMS amplitude on the other (Figure 1D), or a time-series plot showing the RMS signals of both muscles over the previous 30 s. To ensure real-time visualization, the feedback was optimized for low-latency streaming by using memory-mapped arrays and a sliding RMS computation window of 300 ms with continuous buffer updates. The computational delay introduced by the software processing pipeline was less than 20 ms. However, the 300 ms RMS window inherently introduced a causal delay, as the RMS amplitude at any given time reflects the muscle activity over the preceding 300 ms. This small but non-negligible delay was consistent across the session and was found to be brief enough to allow participants to fine-tune their activation strategies in near real-time. The feedback software dynamically scaled each channel to emphasize relative differences between muscles rather than absolute magnitudes.

EMG signals were recorded using the OT Bioelettronica Quattrocento system, sampled at 10,240 Hz and band-pass filtered between 10–5,000 Hz.

### Data Processing

Initial analysis focused on evaluating RMS signals from individual intramuscular channels to assess whether participants achieved selective activation of VM and VL. Trial selection was based exclusively on visual inspection of RMS amplitudes, without the use of quantitative metrics such as correlation coefficients. Amplitudes were reviewed offline to identify recordings that exhibited clear, alternating activation of the two muscles. Two trials from each participant were selected for further analysis, each containing both a VM-focused and a VL-focused task segment presented in immediate succession. To ensure sufficient and stable motor unit activity for reliable decomposition, each selected segment had a minimum duration of 3 s.

Each muscle was recorded with three intramuscular electrodes and one HDsEMG electrode. In the VL, all channels displayed similar activation patterns, so the channel with the best signal quality was chosen for analysis. In contrast, the proximal VM electrode often exhibited distinct activity compared with distal sites of the VM and the whole VL. Therefore, we prioritized the proximal site of the VM for analyses to capture potential spatial differences in neural input.

Selected data segments were subjected to offline motor unit decomposition using **EMGLab**, a validated manual decomposition toolbox, running in MATLAB (Version R2024b, The MathWorks, Inc., Natick, Massachusetts, United States). Motor unit spike trains were manually reviewed and edited to ensure accurate identification of discharge times and assigned to the respective muscle. Spike rasters were inspected to determine whether individual motor units were active during VM- or VL-focused tasks. Discharge rates and cumulative spike trains were computed and compared between VM- and VL-focused strategies (Figure 1E).

To assess decomposition quality, reconstructed EMG signals were generated from identified spike trains and compared with the original recordings (Figure 2A). The residual EMG, defined as the difference between original and reconstructed signals, was expressed as a percentage of total EMG power. Only data segments with residual EMG power less than 30% were retained for further analysis (Luca et al., 2015; Negro, Muceli et al., 2016), indicating that the extracted motor units accounted for most of the signal variance.

**Figure 2.**
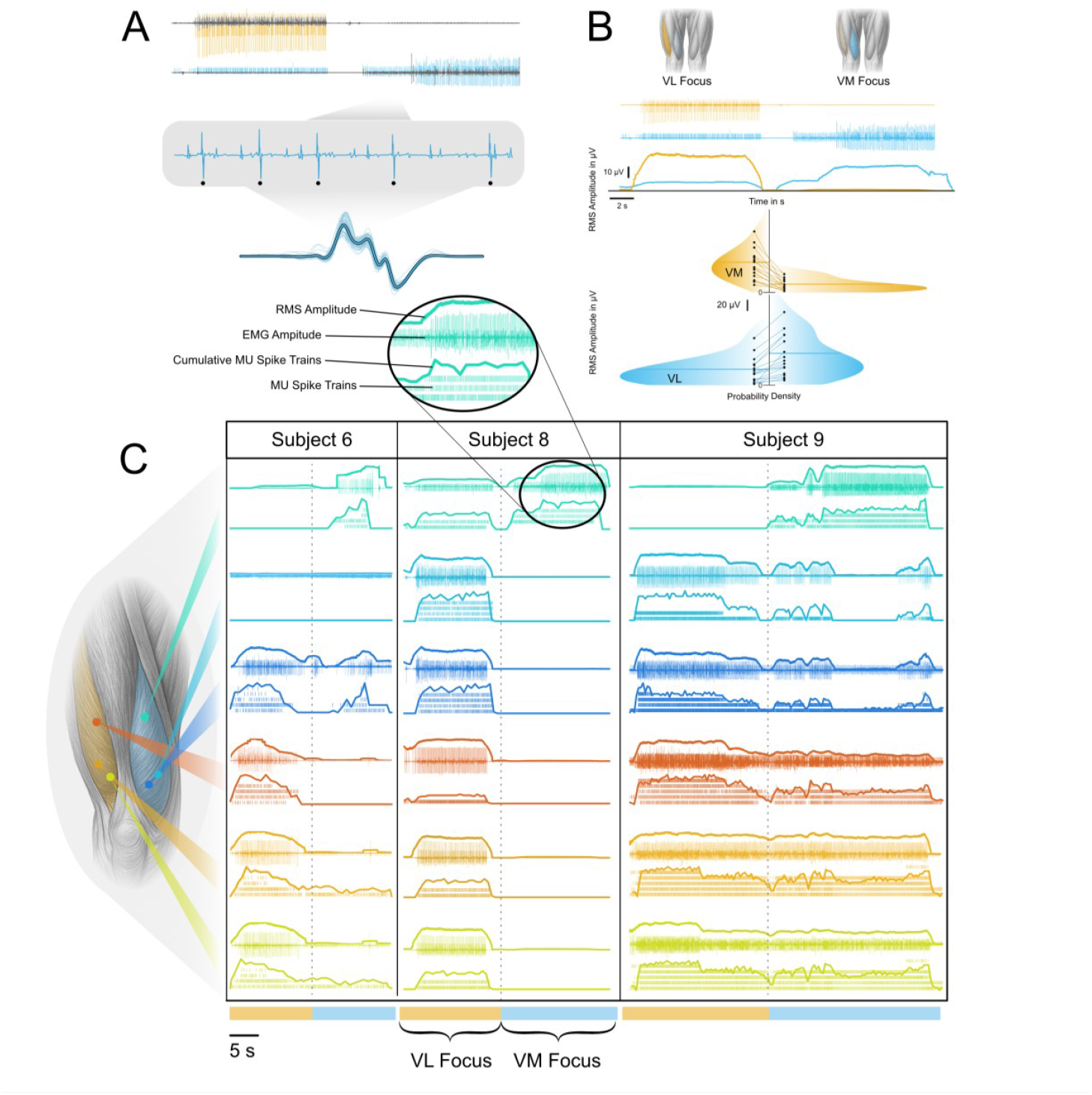
**(A) Validation and visualization of intramuscular EMG decomposition** The upper part of the figure shows intramuscular EMG signals from VM (blue) and VL (yellow) channels, overlaid with the residual EMG after decomposition (gray), highlighting the portion of the signal not explained by the identified motor unit spike trains. A zoomed section of the EMG trace displays individual motor unit action potentials, with five discharges of the same motor unit highlighted to visualize their consistent waveform. The lower part of the figure shows the spike-triggered average (STA) for the selected motor unit with the identified waveforms are superimposed, and the resulting average motor unit action potential shape is indicated by the thick line. **(B) Task-specific modulation of EMG and RMS signals in VM and VL** The top row illustrates the instructed task. VM and VL are alternately highlighted to indicate periods where participants were instructed to focus on selectively activating either muscle. Below, intramuscular EMG signals from VM (blue) and VL (yellow) are shown over time, followed by the corresponding RMS amplitude traces. The bottom panels summarize group-level results from all participants (N = 9), displaying the probability density distributions of RMS amplitudes for VM and VL separately in the corresponding colors. The distributions are shown for VM-focused and VL-focused tasks for each muscle. **(C) Overview of multi-channel and multi-subject motor unit recruitment** EMG activity across all intramuscular channels for three representative participants. The colored traces are matched to the electrode locations. The EMG traces (overlaid with the RMS envelopes) are shown for each participant during both task conditions. Below the raster plots visualize the motor unit spike trains, with each row representing a unique unit. Cumulative discharge rates are overlaid on top to reflect overall modulation of recruitment across tasks.

### Statistical Analysis

All statistical analyses were performed in MATLAB. Differences in RMS amplitude, as well as the number of active motor units, were assessed using Generalized Linear Mixed Models (GLMMs), with Muscle (VM vs. VL) and Task (VM-focused vs. VL-focused) specified as fixed effects and Subject as a random intercept. The proportion of unique motor units was also analyzed using GLMMs with similar fixed and random effects structures. For the analysis of regional differences within VM, pairwise correlation coefficients between RMS signals from proximal and distal channels, as well as between VM and VL channels, were computed for each trial. These correlation values were then compared across conditions using separate GLMMs to test for significant differences between muscle regions. Significance was set at p < 0.05.

## Results

This study aimed to determine whether humans can volitionally dissociate neural inputs to two synergistic muscles, the VM and VL, during isometric contractions. To address this question, we first examined global EMG activity using EMG amplitude analysis to assess gross differences in muscle activation across tasks. Of the ten participants tested, nine were able to achieve a clear separation of VM and VL activity based on the intramuscular EMG amplitude signals. We then performed motor unit decomposition to investigate whether distinct sets of motor units were selectively recruited when the individuals successfully controlled VM or VL in the prescribed tasks. Finally, we validated the decomposition quality and assessed the consistency of motor unit activation across trials.

### EMG Amplitude Analysis

As an initial assessment of the ability to activate the VM and VL selectively, we analyzed the EMG amplitude by computing normalized RMS amplitudes of the intramuscular EMG recordings across the different tasks. Subsequently (offline), we extracted two trial segments from every participant that included the sequential control of VM- and VL-focused tasks. In each participant, the channel demonstrating the clearest task-dependent modulation was selected for further comparison. The clearest channel that showed task-dependent modulation was the proximal channel in the VM (7/9 subjects), whereas in VL all channels showed similar modulation profiles for all participants. The other two participants exhibited clear individual control of VM with both proximal and distal channels showing comparable task-dependent modulation.

In summary, most participants (9/10) were able to separate VM activity from the VL and the VL activity from VM. Figure 2B shows EMG and RMS signals from one participant during a single trial in which the task focus alternated between VM and VL. The upper panel illustrates distinct modulation of intramuscular EMG amplitude in both muscles corresponding to the VM- and VL-focused tasks. Two representative instances of selective activation were identified from each participant (n=9) and further analyzed. A Generalized Linear Mixed Model (GLMM) confirmed that average RMS amplitudes differed significantly between muscles and task conditions. There were significant main effects of Muscle (VM vs. VL; p = 0.001) and Task (VM focus vs. VL focus; p < 0.001), as well as a significant Muscle × Task interaction (p = 0.001). Predicted mean RMS values were greatest for VM during VM-focused tasks (55.3 µV) and for VL during VL-focused tasks (53.1 µV). This is evident in the lower part of Figure 2B, where RMS levels between tasks are directly compared across all subjects.

These results demonstrate task-dependent modulation of VM and VL activity across participants and provided a basis for further analyses at the level of individual motor units.

### Motor Unit Recruitment

To verify that the task-related modulation of RMS amplitude was driven by changes in underlying neural activity, we decomposed the intramuscular EMG signals into individual motor unit spike trains. The decomposition was applied to segments from trials previously identified in the RMS analysis as showing strong selective activation of VM or VL.

To confirm the quality of the intramuscular EMG decomposition, we evaluated the residual EMG power remaining after subtracting the decomposed motor units in the reconstructed signal from the original recordings (Figure 2A). Residual power was expressed as a percentage of the total EMG power. Only segments with residual power below 25 % were included in further analyses (Luca et al., 2015; Negro, Muceli et al., 2016). Across all recordings, 18 trial files were initially selected for decomposition (two per participant). Of these, four files could not be successfully decomposed due to either poor signal quality or excessive activation levels. Consequently, only six participants remained with usable pairs of trials for analyses of trial-to-trial consistency. The residual power for the 14 successfully decomposed recordings averaged 18.04 ± 5.8 %.

Figure 3A illustrates a representative example from two trials of one participant. The top row indicates the instructed task segment (VM- or VL-focused task), followed by the EMG traces from VM and VL. The extracted discharge times of decomposed motor units are shown in the lower panel. During the VM-focused task, a distinct subset of motor units was recruited in VM (marked red) with minimal VL activity. This pattern was reversed during the VM-focused task. This recruitment pattern was stable across trials, with the same motor units consistently activated during VM- or VL-focused tasks. All channels during the 3 trials of different participants are displayed in Figure 2C.

**Figure 3.**
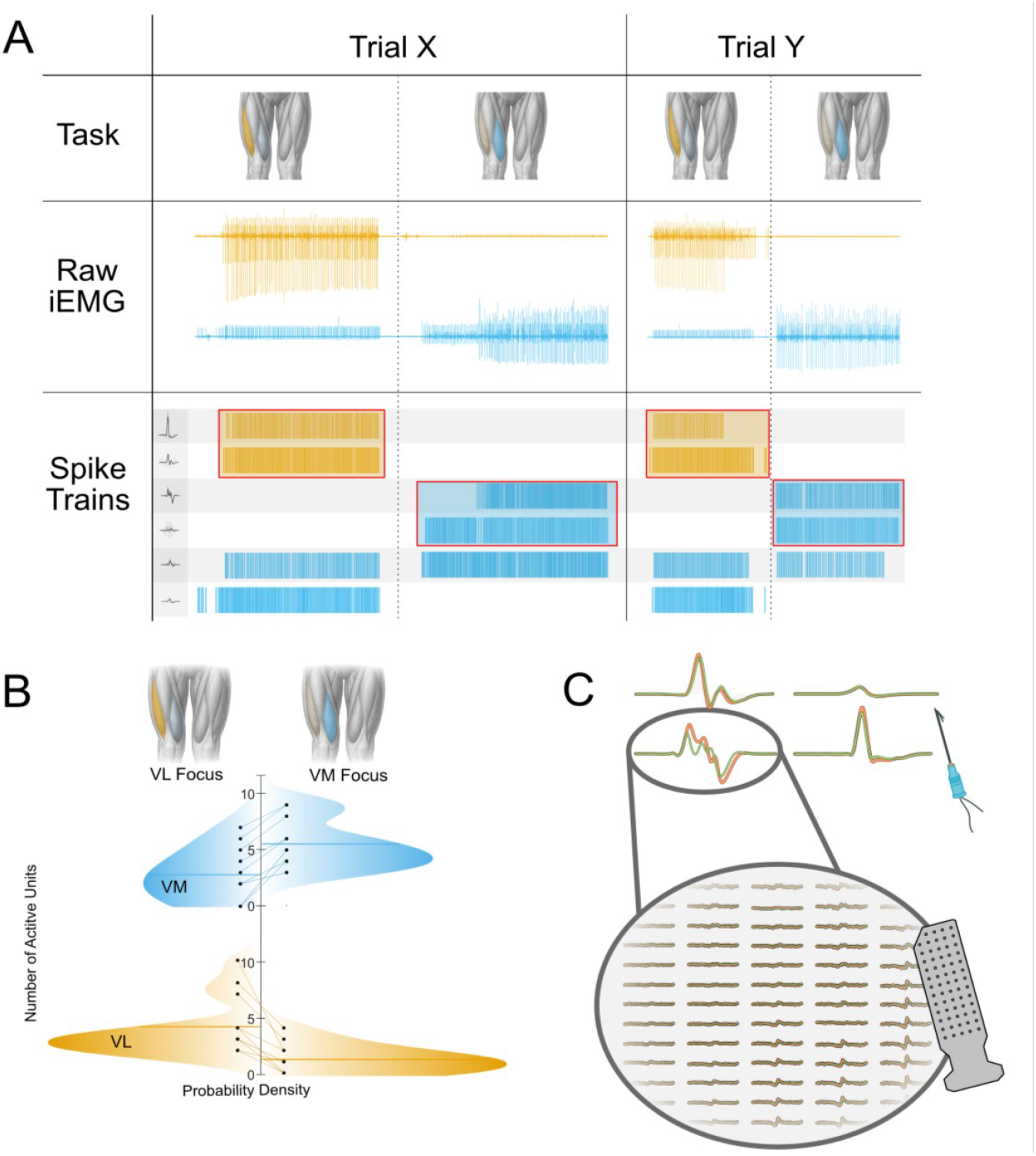
**(A) Comparison of neural drive and motor unit activity across two separate trials in one participant** This compares activation pattern of two trials from the same subject. The top row illustrates the instructed task sequence for each trial, highlighting periods of VL-focused (orange shading) and VM-focused (blue shading) effort. Below, raw intramuscular EMG signals are shown for one channel each from VL and VM, plotted over time and aligned with the task segments. The bottom row displays the corresponding spike trains of all identified motor units, color-coded by muscle (blue for VM, orange for VL), revealing how motor unit recruitment patterns change between VM- and VL-focused segments and how consistent neural drive patterns are across separate trials. **(B) Distribution of active motor units across tasks** Probability density plots are displayed summarizing the number of active motor units identified across all trials used for analysis. For each participant, two trials per task were selected based on clear task-dependent modulation observed in RMS signals. Separate distributions are shown for VM-focused and VL-focused tasks, with data color-coded by muscle (blue for VM, yellow for VL). The y-axis represents the number of active motor units per trial, while the x-axis indicates the estimated probability density. Horizonal lines mark the mean number of active units for each distribution. **(C) MUAP Tracking using Intramuscular and Surface Electrode Recordings** The figure shows the approach used to track individual motor units across trials by combining intramuscular and surface EMG signals. The top panel displays examples of motor unit action potentials (MUAPs) obtained via spike-triggered averaging (STA) of the intramuscular EMG recordings, illustrating the characteristic waveform shapes of individual units. The bottom panel presents the corresponding STA maps projected onto the high-density surface EMG (HDsEMG) grid, visualized as spatial distributions of MUAP amplitude across electrodes. These spatial “footprints” serve as unique signatures, enabling identification of the same motor units across different trials based on both their temporal firing patterns and spatial characteristics.

A GLMM analysis showed a significant interaction between muscle and task (p < 0.001), demonstrating that the number of recruited motor units varied across tasks. On average, the VM exhibited the greatest number of active motor units during the VM-focused task (5.5 ± 2.2 units), whereas during, the number of active VM units dropped to 2.8 ± 2.3 units during the VL-focused task. In contrast, the VL displayed greater recruitment during VL-focused tasks, averaging 4.2 ± 2.6 units, which decreased to 1.33 ± 1.2 units during VM-focused tasks. Although participants were able to modulate motor unit recruitment across tasks, full derecruitment of all motor units was only seen in two participants. These differences are visualized in Figure 3B.

We also analyzed the proportion of units that were uniquely recruited during each task. On average, 71.2% of VL motor units were uniquely recruited during VL-focused tasks, whereas 60.8% of VM motor units were unique to VM-focused tasks. However, this difference was not statistically significant (p = 0.259), indicating that both muscles exhibited comparable levels of task-specific neural recruitment. Although participants were able to selectively activate different sets of motor units, the amount of separation between VM and VL was not consistent.

To determine whether the same motor units exhibited consistent behavior across repeated trials, we tracked individual units using both intramuscular and surface EMG signals (Figure 3C). Spike-triggered averages (STAs) of the intramuscular EMG were computed for each unit, and pairwise correlations were calculated between STAs from different trials. Units were considered matched if they showed the highest correlation across trials and exceeded a threshold of r > 0.9, ensuring temporal consistency in discharge times. In addition, the identity of tracked units was validated spatially by projecting the spike trains onto concurrently recorded HDsEMG grids to generate MUAP maps. High spatial correlation (r < 0.85) of MUAP maps across trials confirmed that matched motor units maintained consistent spatial signatures.

This analysis was performed in the six participants who provided two high-quality decomposable trials. Within these six individuals, we also examined motor units that were identified in both trials to assess the stability of the recruitment activity. Of the motor units tracked across trials, 25 out of 26 VM units (96.2%) demonstrated consistent activity, being either exclusively active during one task or consistently shared across tasks in both trials. Similarly, 15 out of 18 tracked units (83.3%) in VL exhibited stable task-specific activity. This discrimination of motor unit activity was evident in all motor units (Figure 3A).

### Vastus Medialis Compartment Analysis

Inspection of the intramuscular EMG recordings from both VM and VL tasks revealed a consistent profile across all participants. The distal VM and VL channels typically showed concurrent modulation, whereas the proximal VM channel displayed a distinctly different profile (Figure 4A). neuromuscular compartments. We examined correlations between EMG signals (40-100 s) recorded from proximal and distal intramuscular channels across all 18 analyzed trials of the nine participants included in the study. Correlation coefficients were computed between the relevant channels across the entire signal for each trial. Figure 5A presents an example trial in which the RMS signals from the proximal VM, distal VM, and distal VL are displayed together. In this example, the distal VM and distal VL signals were similar, showing concurrent modulation during the task (r=0.83). In contrast, the proximal VM channel demonstrates a distinctly different profile (0.11) compared with the distal VM channel. Data from all channels across trials are displayed in Figure 4A, which shows clear separation of the activity between these two clusters. The proximal site of VM muscle to receive a different neural drive than the other channels implanted in VM and VL. This pattern was clearly observable in 6/9 participants, with the proximal part of the VM showing uncorrelated EMG amplitude signal relative to VL and distal VM.

**Figure 4.**
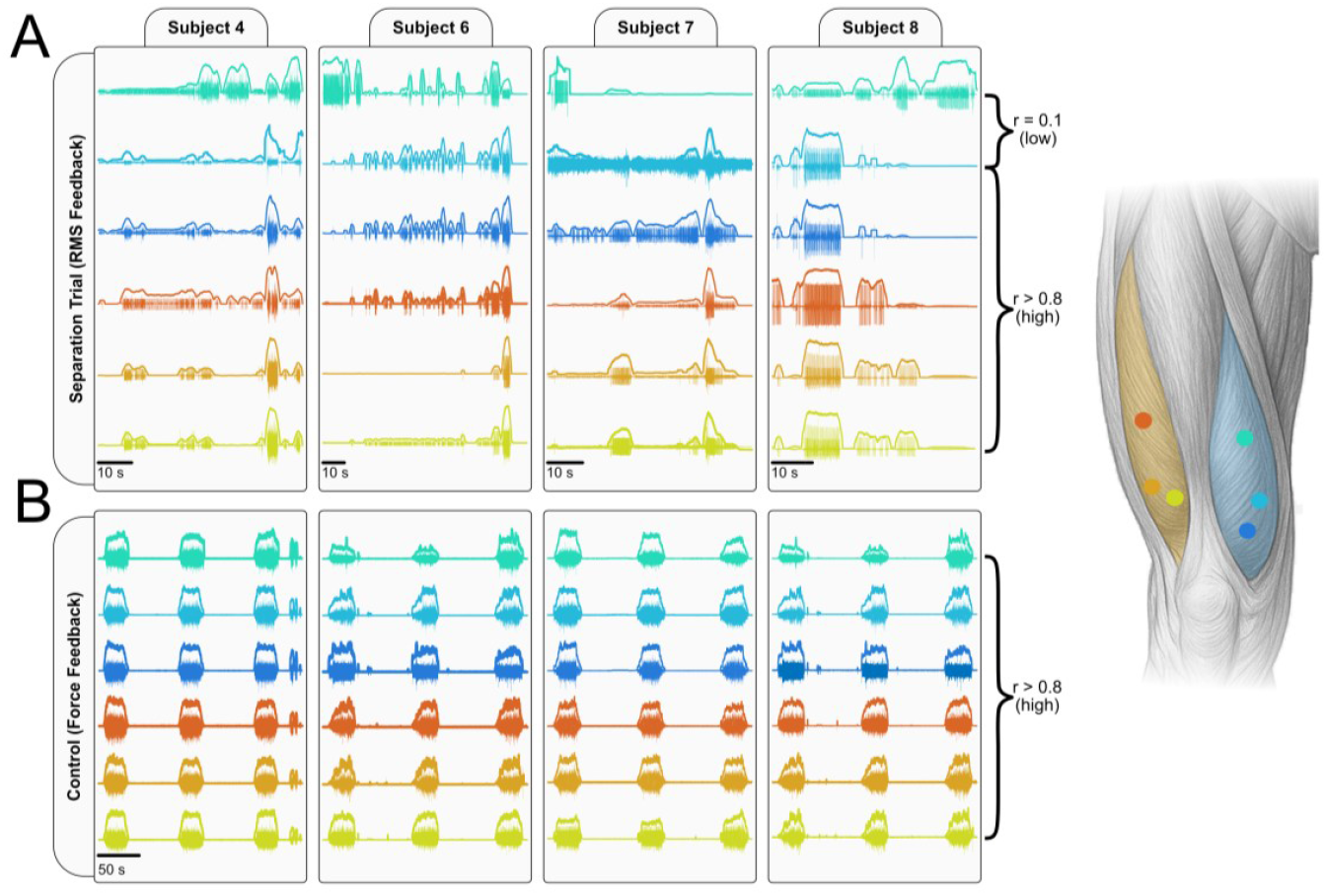
**Overview of intramuscular EMG activity in vastus medialis and vastus lateralis across four subjects** This panel presents representative intramuscular EMG recordings from both the feedback condition (RMS-guided selective activation) and the control condition (force feedback) across four subjects. EMG signals from all implanted channels (color coded) are displayed for each subject. EMG traces are overlaid with their respected RMS amplitude and highlight differences in concurrent modulation across channels. All channels show tightly coupled in the control condition, whereas proximal VM channels often exhibit independent modulation relative to distal VM and VL in the feedback condition. On the right, an anatomical schematic of the anterior thigh illustrates the locations of all intramuscular electrodes. Colored dots correspond to specific EMG channels plotted on the left, enabling anatomical interpretation of the spatial activation differences observed across conditions.

**Figure 5.**
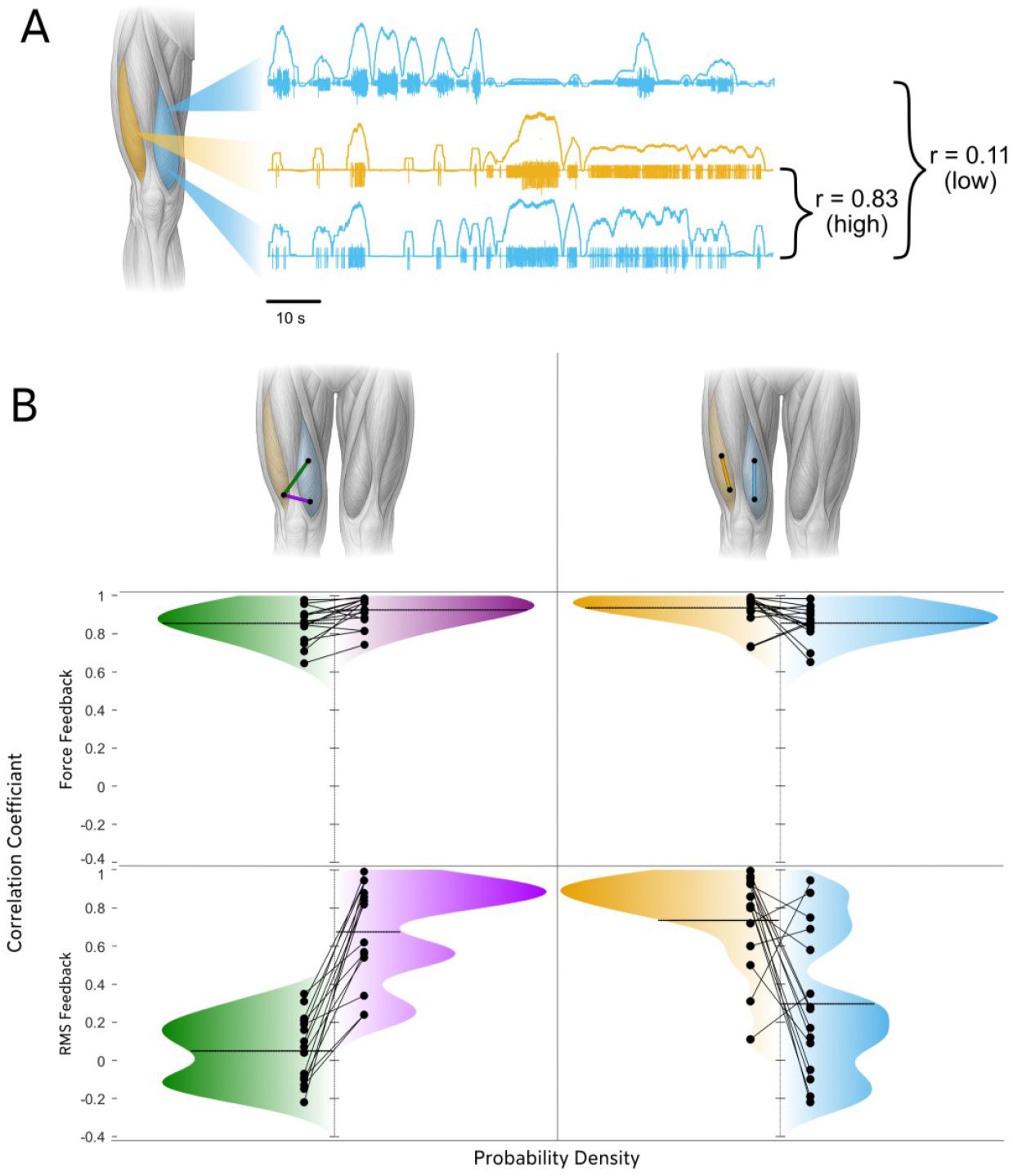
**(A) Visualization of regional EMG recording and correlation analysis** A schematic of the thigh shows the locations of VL (yellow) and VM (blue). Intramuscular EMG recordings acquired concurrently from different muscles and regions (distal/proximal) of the muscle are displayed next to the muscle. Above each EMG trace, the RMS signals are displayed to illustrate the amplitude modulation over time. Correlation coefficients demonstrate a high correlation (r = 0.83) between VL and distal VM signals, but a lower correlation (r = 0.11) between distal and proximal regions of VM. **(B) Distributions of RMS correlation coefficients across muscle regions** This figure illustrates the distributions of correlation coefficients computed from RMS signals recorded across different quadriceps regions during both the EMG feedback condition and the ramp contractions (force feedback). Each subplot displays a continuous probability density plot, where the vertical axis represents correlation values and the horizontal axis indicates probability density. The top row presents the distributions from the ramp contractions (15% and 30% MVC ramp contractions with force feedback), whereas the bottom row displays the corresponding distributions from the EMG feedback task. Each column shows a specific pairwise comparison: (1) distal VL vs. proximal VM, (2) distal VM vs. distal VL, (3) distal vs. proximal VL, and (4) distal vs. proximal VM.

To quantify these relations across all participants and trials, we computed pairwise correlation coefficients for each muscle pair. As summarized in Figure 5B, correlations between the proximal and distal channels of VL were consistently high, with a mean correlation of 0.76 ± 0.08. In contrast, correlations between the distal and proximal regions of VM were significantly lower, averaging only 0.39 ± 0.12. A GLMM confirmed this difference (p = 0.0022), indicating that the proximal and distal regions of VM did not function as a single functional unit. Moreover, the cross-muscle correlation between distal VM and distal VL was 0.68 ± 0.06, whereas it was 0.12 ± 0.09 between proximal VM versus distal VL. The GLMM revealed this difference to be highly significant (p < 0.001).

To assess whether this compartmentalization was specific to the feedback-driven condition, we examined standard isometric ramp contractions with force feedback instead of EMG feedback. In contrast to the EMG feedback condition, EMG signals were strongly correlated during the ramp contractions (Figure 4B) with mean correlation coefficient was 0.85 ± 0.097for proximal VM vs distal VL, 0.92 ± 0.074 for distal VM vs distal VL, 0.86 ± 0.089 for proximal VM vs distal VM, and 0.93 ± 0.082 for proximal VL vs distal VL.

### Force Applied to the Dynamometer

The lower leg was attached to the dynamometer to minimize movement in the side-to-side (x), upward-downward (y-axis) and forward-backward (z) directions, (Figure 6A). Several participants (6 of 9) reported that selectively activating the vastus medialis (VM) or vastus lateralis (VL) was easier when applying downward pressure through the foot, which was accommodated by the addition of a small foot platform.

**Figure 6.**
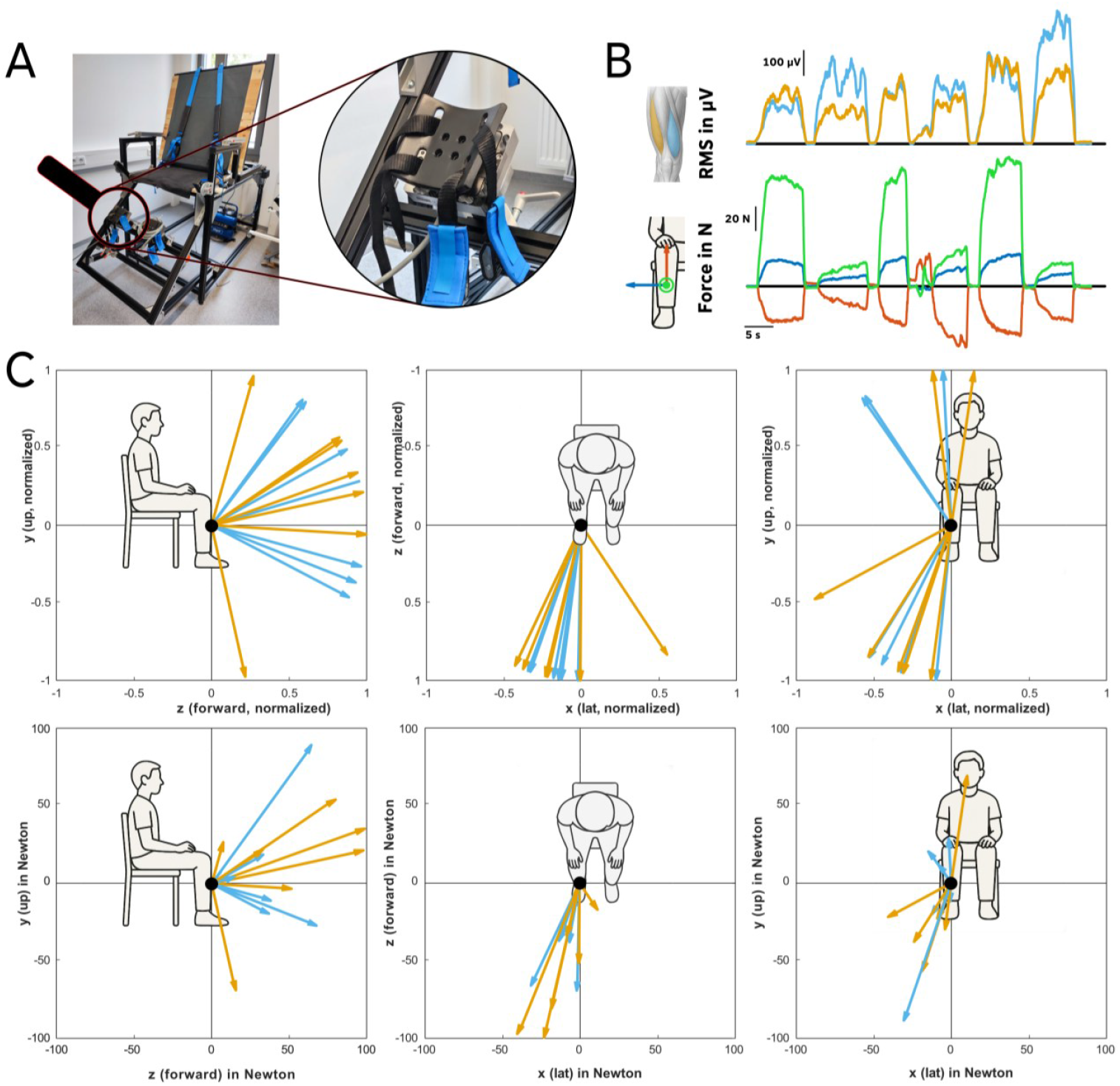
**(A) Photograph of the dynamometer with the integrated three-dimensional force sensor** The lower leg was strapped to the lever arm to restrict movement in the side-to-side (x), upward-downward (y) and forward-backward (z) directions. **(B) *Example recording*** The upper plot shows the RMS of intramuscular EMG signals from a representative trial. The lower plot displays the corresponding three-dimensional force signals recorded by the force sensor: green = forward-backward, blue = side-to-side, and red = upward=downward. **(C) Force vector visualization across tasks** Two-dimensional projections of force vectors recorded during VM- and VL-focused conditions. Each arrow represents one exemplar vector per participant (blue = VM focus, orange = VL focus). The top row shows normalized unit vectors projected onto the x–y (lateral–up), x– z (lateral–forward), and y–z (up–forward) planes, highlighting task-dependent directional differences. The bottom row shows the same projections with non-normalized vectors (in Newtons), illustrating inter-individual differences in absolute force magnitude.

Across participants, systematic differences in force orientation emerged between the VM- and VL-focused tasks (Figure 6C). For each participant, the net force generated by the lower leg was summarized as a single three-dimensional unit vector, indicating the direction in which the force was applied. The coordinate system was defined such that x corresponds to lateral, y to vertical, and z to forward.

During VM-focused contractions, the average direction pointed mainly upward and forward, with only a small lateral component. Numerically, this corresponded to [0.215, -0.210, 0.954]. In the VL-focused condition, the mean direction was [0.180, -0.296, 0.938], pointing in a similar but slightly different direction. The angle between these two mean directions was 5.3°, indicating a small but measurable difference in how participants oriented their force across tasks.

To make this difference more explicit, we subtracted the VM mean direction vector from the VL mean direction vector (ΔV = V_VL - V_VM). This difference vector illustrates how the orientation of the applied force shifted between tasks. The components of ΔV were Δx = -0.035, Δy = -0.086, and Δz = -0.016, indicating that the largest average change occurred in the vertical (y) direction.

Finally, to determine the most systematic pattern of change across participants, we performed principal component analysis (PCA) on the trial-by-trial difference vectors. The first principal component explained 59 % of the variance and was dominated by the vertical (y) axis, indicating that vertical modulation was the most consistent factor distinguishing VM- and VL-focused tasks, even though the overall difference in mean directions was small. Taken together, these results show that although the global force orientations for VM- and VL-focused contractions were largely similar, systematic task-dependent differences emerged primarily in the vertical direction.

## Discussion

This study investigated whether humans can volitionally dissociate activity between VM and VL, as well as within compartments of the VM, during low-force isometric contractions. By combining intramuscular EMG, high-density surface recordings, and real-time feedback, the findings provide evidence that region-specific modulation within synergistic muscles is possible during a short-term practice intervention

Participants (9/10) were generally able to differentially modulate VM and VL using EMG feedback, but complete dissociation, defined as selective activation of one region with no coactivation of the other, was rarely achieved. Although two individuals demonstrated near-complete task-specific motor unit recruitment in VM or VL, most participants exhibited partial overlap in motor unit activity across tasks. This suggests that volitional dissociation is not universally accessible, and that the neural and anatomical architecture of the quadriceps imposes strong coupling that limits independent control (Laine et al., 2015; Luca and Erim, 1994).

A key finding was the asymmetric flexibility of control across muscle regions. Although the proximal VM often exhibited activation patterns distinct from both distal VM and VL, this distinction was not spontaneous. In standard ramp contractions without EMG feedback, muscle activity was highly correlated (r > 0.8), underscoring that region-specific dissociation only emerged with explicit feedback and focused volitional effort. Even then, selective activation of VL or distal VM was notably more difficult to achieve than proximal VM, indicating directional biases in controllability that may reflect underlying differences in neural input or functional roles. Our correlation analyses confirmed these observations. The sharp contrast between conditions demonstrates that feedback facilitates a mode of neuromuscular control that is otherwise latent, and that task-dependent reorganization of motor unit activity can emerge with brief practice.

These findings are consistent with previous literature suggesting partial independence of VM and VL at the neural level (Del Vecchio et al., 2023; Hug et al., 2023), and extend those insights by offering direct, motor unit-level evidence of region-specific recruitment. Moreover, the proximal VM has been anatomically characterized as a distinct compartment (Gallina et al., 2016; Travnik et al., 1995), and our data are consistent with the notion that this region may be preferentially targeted during specific motor tasks. Selective recruitment of motor units within proximal VM, accompanied by consistent reductions in coactivation with VL, supports the existence of functionally relevant compartmentalization within the vastus medialis (English et al., 1993; Gallina et al., 2016; Travnik et al., 1995).

Although motor unit recruitment strategies were reproducible within individuals across trials, they were not transferable between participants. The specific strategies (e.g., inward rotation, pressure distribution) that enabled successful dissociation were highly individualized. This individuality reinforces the idea that volitional dissociation is not achieved through a single universal motor pattern but rather through self-exploration, and aligns with recent findings of task-specific recruitment in other muscle systems (Cakici et al., 2022; Oßwald et al., 2025; Simpetru et al., 2024). Together, these observations suggest that while the neuromuscular system can establish stable, task-specific strategies within subjects, full dissociation more likely emerges as a learned skill.

Our force-vector analysis further emphasized this individuality. Although average differences suggested slightly larger forward-backward (z) components, PCA revealed that vertical (y-axis) modulation was the most consistent distinguishing factor across participants. This suggests that upward force control plays a key role in enabling selective VM versus VL activation, even though absolute magnitudes of this force was not measured. Importantly, this interpretation must be considered in light of a methodological limitation: the foot rested on a small platform that was not mechanically connected to the dynamometer’s force transducer, likely leading to an underestimation of the absolute vertical force component. Nevertheless, the y-axis contribution was still clearly detectable in the recordings, indicating that upward force played a central role in enabling selective VM versus VL activation.

From a methodological standpoint, our combination of intramuscular decomposition and surface mapping enabled high-resolution assessment of spatial activation patterns. However, decomposition was limited to low-force contractions due to signal overlap at higher intensities, restricting the generalizability of our findings to functional or athletic movements. Nonetheless, the ability to volitionally modulate specific motor unit pools, particularly in the proximal VM, could support targeted neuromuscular retraining in populations with patellofemoral pain or post-surgical quadriceps imbalance. Importantly, however, the need for explicit feedback and guided practice highlights that such control is not spontaneously expressed, and that region-specific interventions may require individualized strategies and long-term engagement.

## Conclusion

Our study provides evidence that the central nervous system possesses some capacity to flexibly modulate neural inputs to the quadriceps, enabling some regional recruitment of distinct sets of motor units. The compartmentalization observed within VM suggests that this muscle is not homogeneously controlled but instead includes regions that can be differentially activated under specific volitional strategies. However, full dissociation between VM and VL, or between VM compartments, was rare. These findings refine our understanding of neuromuscular control in this large group of synergistic muscles and open new avenues for research and clinical intervention aimed at enhancing selective motor control through feedback-guided training and personalized rehabilitation strategies.

## References

Caillet, A. H., Avrillon, S., Kundu, A., Yu, T., Phillips, A. T. M., Modenese, L. and Farina, D. (2023) ‘Larger and Denser: An Optimal Design for Surface Grids of EMG Electrodes to Identify Greater and More Representative Samples of Motor Units’, eNeuro, vol. 10, no. 9.

Cakici, A. L., Osswald, M., Oliveira, D. S. de, Braun, D. I., Simpetru, R. C., Kinfe, T., Eskofier, B. M. and Del Vecchio, A. (2022) ‘A Generalized Framework for the Study of Spinal Motor Neurons Controlling the Human Hand During Dynamic Movements’, Annual International Conference of the IEEE Engineering in Medicine and Biology Society. IEEE Engineering in Medicine and Biology Society. Annual International Conference, vol. 2022, pp. 4115–4118.

d’Avella, A., Portone, A., Fernandez, L. and Lacquaniti, F. (2006) ‘Control of fast-reaching movements by muscle synergy combinations’, The Journal of neuroscience : the official journal of the Society for Neuroscience, no. 30, pp. 7791–7810.

d’Avella, A., Saltiel, P. and Bizzi, E. (2003) ‘Combinations of muscle synergies in the construction of a natural motor behavior’, Nature neuroscience, vol. 6, no. 3, pp. 300–308.

Del Vecchio, A., Marconi Germer, C., Kinfe, T. M., Nuccio, S., Hug, F., Eskofier, B., Farina, D. and Enoka, R. M. (2023) ‘The Forces Generated by Agonist Muscles during Isometric Contractions Arise from Motor Unit Synergies’, The Journal of neuroscience : the official journal of the Society for Neuroscience, vol. 43, no. 16, pp. 2860–2873.

Dernoncourt, F., Avrillon, S., Logtens, T., Cattagni, T., Farina, D. and Hug, F. (2024) Flexible Control of Motor Units: Is the Multidimensionality of Motor Unit Manifolds a Sufficient Condition?

Desmedt, J. E. and Godaux, E. (1978) ‘Ballistic contractions in fast or slow human muscles: Discharge patterns of single motor units’, J Physiol, no. 285, pp. 185–196.

Desmedt JE, G. E. (1977) ‘Fast motor units are not preferentially activated in rapid voluntary contractions in man’, Nature, no. 267, pp. 717–719.

English, A. W., Wolf, S. L. and Segal, R. L. (1993) ‘Compartmentalization of Muscles and Their Motor Nuclei: The Partitioning Hypothesis’, Physical Therapy, no. 73, pp. 857–867.

Farina, D., Merletti, R., Indino, B. and Graven-Nielsen T. (2004) ‘Surface EMG crosstalk evaluated from experimental recordings and simulated signals: reflections on crosstalk interpretation, quantification and reduction.’, Methods Inf Med, no. 43, pp. 30–35.

Gallina, A., Ivanova, T. D. and Garland, S. J. (2016) ‘Regional activation within the vastus medialis in stimulated and voluntary contractions’, Journal of applied physiology (Bethesda, Md.: 1985), vol. 121, no. 2, pp. 466–474.

Germer, C. M., Farina, D., Elias, L. A., Nuccio, S., Hug, F. and Del Vecchio, A. (2021) ‘Surface EMG cross talk quantified at the motor unit population level for muscles of the hand, thigh, and calf’, Journal of applied physiology (Bethesda, Md. : 1985), vol. 131, no. 2, pp. 808–820.

Harris, A. J., Duxson, M. J., Butler, J. E., Hodges, P. W., Taylor, J. L. and Gandevia, S. C. (2005) ‘Muscle fiber and motor unit behavior in the longest human skeletal muscle’, The Journal of neuroscience : the official journal of the Society for Neuroscience, vol. 25, no. 37, pp. 8528–8533.

Heckmann, C. J., Gorassini, M. A. and Bennett, D. J. (2005) ‘Persistent inward currents in motoneuron dendrites: implications for motor output’, Muscle & nerve, vol. 31, no. 2, pp. 135–156.

Hug, F., Avrillon, S., Sarcher, A., Del Vecchio, A. and Farina, D. (2023) ‘Networks of common inputs to motor neurons of the lower limb reveal neural synergies that only partly overlap with muscle innervation’, J Physiol, no. 601.15, pp. 3201–3219.

Ivanenko, Y. P., Poppele, R. E. and Lacquaniti, F. (2004) ‘Five basic muscle activation patterns account for muscle activity during human locomotion’, The Journal of physiology, vol. 556, Pt 1, pp. 267–282.

Laine, C. M., Martinez-Valdes, E., Falla, D., Mayer, F. and Farina, D. (2015) ‘Motor Neuron Pools of Synergistic Thigh Muscles Share Most of Their Synaptic Input’, The Journal of neuroscience : the official journal of the Society for Neuroscience, vol. 35, no. 35, pp. 12207–12216.

Levine, J., Avrillon, S., Farina, D., Hug, F. and Pons, J. L. (2022) Two motor neuron synergies, invariant across ankle joint angles, activate the triceps surae during plantarflexion.

Luca, C. J. de and Erim, Z. (1994) ‘Common drive of motor units in regulation of muscle force’, Trends Neurosci, no. 17, pp. 299–305.

Luca, C. J. de, Nawab, S. H. and Kline, J. C. (2015) ‘Clarification of methods used to validate surface EMG decomposition algorithms as described by Farina et al. (2014)’, Journal of applied physiology (Bethesda, Md. : 1985), vol. 118, no. 8, p. 1084.

Madarshahian, S., Letizi, J. and Latash, M. L. (2021) ‘Synergic control of a single muscle: The example of flexor digitorum superficialis’, The Journal of physiology, vol. 599, no. 4, pp. 1261–1279.

Marshall, N. J., Glaser, J. I., Trautmann, E. M., Amematsro, E. A., Perkins, S. M., Shadlen, M. N., Abbott, L. F., Cunningham, J. P. and Churchland, M. M. (2022) ‘Flexible neural control of motor units’, Nature neuroscience, vol. 25, no. 11, pp. 1492–1504.

Mellor, R. and Hodges, P. (2005) ‘Motor unit synchronization between medial and lateral vasti muscles’, Clinical neurophysiology : official journal of the International Federation of Clinical Neurophysiology, vol. 116, no. 7, pp. 1585–1595.

Milner-Brown, H. S., Stein, R. B. and Yemm, R. (1973) ‘The contractile properties of human motor units during voluntary isometrix contractions’, J Physiol, no. 228, pp. 285–306.

Negro, F., Muceli, S., Castronovo, A. M., Holobar, A. and Farina, D. (2016) ‘Multi-channel intramuscular and surface EMG decomposition by convolutive blind source separation’, Journal of neural engineering, vol. 13, no. 2, p. 26027.

Negro, F., Yavuz, U. Ş. and Farina, D. (2016) ‘The human motor neuron pools receive a dominant slow-varying common synaptic input’, The Journal of physiology, vol. 594, no. 19, pp. 5491–5505.

Oßwald, M., Cakici, A. L., Souza de Oliveira, D., Braun, D. I., Farina, D. and Del Vecchio, A. (2025) ‘Task-specific motor units in the extrinsic hand muscles control single- and multidigit tasks of the human hand’, Journal of applied physiology (Bethesda, Md. : 1985), vol. 138, no. 5, pp. 1187–1200.

Powers, C. M. (2000) ‘The Influence of Vastus Muscle Activity in Subjects With and Without Patellofemoral Pain’, Physical Therapy, Volume 80, Issue 10, pp. 956–964.

Rossato, J., Avrillon, S., Tucker, K., Farina, D. and Hug, F. (2024) ‘The Volitional Control of Individual Motor Units Is Constrained within Low-Dimensional Neural Manifolds by Common Inputs’, The Journal of neuroscience : the official journal of the Society for Neuroscience, vol. 44, no. 34.

Sandercock, T. G., Wei, Q., Dhaher, Y. Y., Pai, D. K. and Tresch, M. C. (2018) ‘Vastus lateralis and vastus medialis produce distinct mediolateral forces on the patella but similar forces on the tibia in the rat’, Journal of biomechanics, vol. 81, pp. 45–51.

Simpetru, R. C., Souza de Oliveira, D., Ponfick, M. and Del Vecchio, A. (2024) ‘Identification of Spared and Proportionally Controllable Hand Motor Dimensions in Motor Complete Spinal Cord Injuries Using Latent Manifold Analysis’, IEEE transactions on neural systems and rehabilitation engineering : a publication of the IEEE Engineering in Medicine and Biology Society, vol. 32, pp. 3741–3750.

Travnik, L., Pernus, F. and Erzen, I. (1995) ‘Histochemical and morphometric characteristics of the normal human vastus medialis longus and vastus medialis obliquus muscles’, Journal of Anatomy, no. 187, pp. 403–411.

Tresch, M. C., Cheung, V. C. K. and d’Avella, A. (2006) ‘Matrix factorization algorithms for the identification of muscle synergies: evaluation on simulated and experimental data sets’, Journal of neurophysiology, vol. 95, no. 4, pp. 2199–2212.

Weinman, L. E., Del Vecchio, A., Mazzo, M. R. and Enoka, R. M. (2024) ‘Motor unit modes in the calf muscles during a submaximal isometric contraction are changed by brief stretches’, The Journal of physiology, vol. 602, no. 7, pp. 1385–1404.

